# Investigating the biomechanical properties of streptococcal polysaccharide capsules using atomic force microscopy

**DOI:** 10.1101/723841

**Authors:** H Marshall, S Aguayo, M Kilian, FC Petersen, L Bozec, JS Brown

**Author notes:** Corresponding author. Mailing address: Centre for Inflammation and Tissue Repair, Department of Medicine, Royal Free and University College Medical School, Rayne Institute, 5 University Street, London WC1E 6JF, United Kingdom. Joint 1^st^ authors (H Marshall, S Aguayo). Joint senior authors (L Bozec, JS Brown).

## Abstract

In common with many bacterial pathogens, *Streptococcus pneumoniae* has a polysaccharide capsule, which facilitates immune evasion and is a key virulence determinant. However, recent data has shown that the closely related *Streptococcus mitis* can also express polysaccharide capsules including those with an identical chemical structure to *S. pneumoniae* capsular serotypes. We have used atomic force microscopy (AFM) techniques to investigate the biophysical properties of *S. mitis* and *S. pneumoniae* strains expressing the same capsular serotypes that might relate to their differences in virulence potential. When comparing *S. mitis* and *S. pneumoniae* strains with identical capsule serotypes *S. mitis* strains were more susceptible to neutrophil killing and imaging using electron microscopy and AFM demonstrated significant morphological differences. Force-volume mapping using AFM showed distinct force-curve profiles for the centre and edge areas of encapsulated *S. pneumoniae* and *S. mitis* strains. This “edge effect” was not observed in the unencapsulated streptococcal strains and in an unencapsulated *Staphylococcus aureus* strain, and therefore was a direct representation of the mechanical properties of the bacterial capsule. When two strains of *S. mitis* and *S. pneumoniae* expressed an identical capsular serotype, they presented also similar biomechanical characteristics. This would infer a potential relationship between capsule biochemistry and nanomechanics, independent of the bacterial strains. Overall, AFM was an effective tool to explore the biophysical properties of bacterial capsules of living bacteria by reproducibly quantifying the elastic and adhesive properties of bacterial cell surfaces. Using AFM to investigate capsule differences over a wider range of strains and capsular serotypes of streptococci and correlate the data with phenotypic differences will elucidate how the biophysical properties of the capsule can influence its biological role during infection.

## Introduction

*Streptococcus pneumoniae* often colonises the human nasopharynx, and is also one of the most common causes of death due to a microorganism, causing a high proportion of global cases of pneumonia, septicaemia and meningitis (Fitzwater et al., 2012, Herman-Bausier and Dufrêne, 2018). *Streptococcus mitis* is the closest genetic relative of *S. pneumoniae* and is a respiratory tract commensal, although more often found in the oropharynx rather than the nasopharynx (Frandsen et al., 1991, Wan et al., 2001). Despite their close genetic relationship, *S. mitis* has a weak virulence potential compared to *S. pneumoniae*. Identifying why these species vary in their ability to cause invasive disease in humans would help characterise why some upper respiratory tract colonisers are also common pathogens whereas most have only weak virulence.

In most strains, the *S. pneumoniae* surface is covered in a polysaccharide capsule consisting of chains of repeating oligosaccharide units. The monosaccharide content, their chemical linkages, and the presence or absence of side-chains creates considerable diversity in capsular structure and antigenicity that is divided into 95+ capsular serotypes (Bentley et al., 2006). The capsule is a well-recognised *S. pneumoniae* virulence determinant required for bacterial evasion of complement- and antibody-mediated phagocytosis (Camberlein et al., 2015, Hyams et al., 2010a). Capsule serotypes differ markedly in their ability to cause invasive infection, with resistance to phagocytosis and susceptibility to complement deposition also varying between serotype and correlating with invasive potential (Hyams et al., 2010a, Hyams et al., 2013). Unencapsulated strains of *S. pneumoniae* do not cause septicaemia or meningitis, and are markedly attenuated in their ability to cause invasive disease in animal models of infection (Briles et al., 1992, Morona et al., 2004, Morona et al., 2006, Rukke et al., 2012), leading to the assumption that *S. mitis* lacked virulence as it does not possess a surface capsule. However, although expression of the *S. pneumoniae* serotype 4 capsule in an *S. mitis* strain improved resistance to complement and phagocytosis this did not make the resulting *S. mitis* TIGR4_*cps*_ strain virulent in mouse models. Furthermore, genome sequencing has shown a proportion of *S. mitis* strains contain a complete *cps* genetic locus arranged almost identically to the *cps* locus of *S. pneumoniae*, and serology and microscopy have confirmed some *S. mitis* strains are surrounded by a capsule (Rukke et al., 2012, Skov Sørensen et al., 2016b). Although the gene content within *S. mitis cps* loci often differs from those in identified *S. pneumoniae* strains, suggesting the capsule has a different biochemical structure, recently *S. mitis cps* loci have been identified that have a gene content that is highly similar to the *cps* loci of some *S. pneumoniae* capsular serotypes (Lessa et al., 2018, Skov Sørensen et al., 2016b, Kilian et al., 2008). These data suggest that through genetic recombination *S. mitis* and *S. pneumoniae* strains can acquire biochemically identical capsule serotypes (Kilian et al., 2014) but raise the question whether these capsules could have other functional differences (e.g. relative levels of expression) that may still contribute to differences in virulence between *S. mitis* and *S. pneumoniae*.

There are extensive epidemiological data on the effects of *S. pneumoniae* capsular serotypes on disease phenotypes, and these have been partially correlated to *in vitro* measures of virulence such as resistance to complement and phagocytosis (Hyams et al., 2011, Weinberger et al., 2009). Increased capsule thickness in opaque phase variants of *S. pneumoniae* compared to transparent phase variants is associated with greater resistance to complement and phagocytosis (Hyams et al., 2010a), and variations in capsule thickness between serotypes has also been correlated with resistance to non-opsonic phagocytosis (Weinberger et al., 2009). These data would suggest that the capsule simply prevents host proteins attaching to the bacterial cell surface. In contrast, we have previously found that strain resistance to complement and neutrophil phagocytosis correlates strongly with the degree of binding of the host protein complement inhibitor factor H to the *S. pneumoniae* subcapsular cell wall protein PspC (Hyams et al., 2013), indicating a more nuanced effect of capsule on inhibiting host immunity than simply blocking access to the bacterial surface by host proteins. How the chemical structure of the capsule affects interactions between host and bacterial molecules is not known, and investigating this will require new methodologies that can measure the physical effects of the capsule on host interactions. Atomic force microscopy (AFM) has proven to be a reliable tool to image and characterise the biomechanical properties of a wide range of bacterial cells under physiological environment conditions (Herman-Bausier and Dufrêne, 2018). As no invasive sample preparation is required for AFM (when compared to other types of microscopy techniques), it is possible to immobilise bacteria under buffered conditions and obtain high-resolution imaging of viable cells (Braga and Ricci, 2011). Recently, AFM has been used to characterise the adhesion of cells to substrates at the single-cell and single-molecule levels (Beaussart et al., 2014), and Wang *et al*. utilised AFM to study the mechanical behaviour of *Klebsiella pneumoniae* strains. They found that the presence of type 3 fimbriae maintained fluidity of the polysaccharide capsule and that this facilitated adhesion to surfaces (Wang et al., 2015). The same group also demonstrated that the *K. pneumoniae* capsule altered its elasticity in response to changes in turgor pressure by absorbing counterions, so reducing the overall net charge along the capsule polysaccharide chains and protecting the bacterial cell against osmotic stress (Wang et al., 2013). Su *et al.*, also used AFM to characterise the structure of the polysaccharide capsule of *Zunongwangia profunda* SM-A87 (Su et al., 2012), while Stukalov *et al.* characterised the capsule of four different gram-negative bacterial strains by utilising both AFM and transmission electron microscopy (TEM) (Stukalov et al., 2008). While TEM allowed visualisation of the capsule for some strains, AFM was able to detect the presence of capsule on all the encapsulated strains studied (Stukalov et al., 2008). These publications confirm that AFM can be used to evaluate the mechanical properties of encapsulated and unencapsulated bacteria and promises to be a good approach for investigating the physical differences related to the capsule between bacterial strains or species (Dufrêne et al., 2017).

In this study, the phenotypic characteristics (structure, mechanical properties, resistance to neutrophil-mediated opsonophagocytosis and killing) of different capsular serotypes associated with *S. mitis* and *S. pneumoniae* strains were investigated and complemented with nanometrology measurement. The overall aims were to investigate whether there are physical differences between *S. mitis* and *S. pneumoniae* strains expressing the same capsular serotypes that might relate to differences in their virulence potential.

## Materials and Methods

### Bacterial strains and culture conditions

The strains of *S. pneumoniae* and *S. mitis* used in this study are listed in Table 1, and have all been described previously (Camberlein et al., 2015, Skov Sørensen et al., 2016b, Spoor et al., 2015). The *Staphylococcus aureus* strains used were kind gifts from R. Fitzgerald, University of Edinburgh. Bacteria were cultured at 37°C in 5% CO2 in air on Colombia blood agar plates supplemented with 5% defibrinated horse blood or in Todd-Hewitt broth supplemented with 0.5% yeast extract (THY). Growth in liquid medium was assessed by optical density. Bacterial cultures were grown to approximately mid-log phase (0.4-0.5 at OD580nm, approximately 1×108 CFU/ml) and stored as single use aliquots at -80°C.

**Table 1:** Bacterial strains used for this study.

| Species | Strain | Serotype |
| --- | --- | --- |
| *S. pneumoniae* | TIGR4 | 4 |
|  | SK618 | 19C |
|  | SK1095/39 | 36 |
|  | SK1442 | 45 |
| | TIGR4Δcps | 4 |
| *S. mitis* | NCTC12261 (SK142) | unknown |
| | SK142 Δcps | - cps |
|  | SK142 TIGR4<sub>cps</sub> | 4 |
|  | SK564 | 19C |
|  | SK1126 | 36 |
|  | CCUG62644 | 45 |
|  | NCTC10712 (SK113) | - cps |
|  | SK1080 | -cps |
| *S. aureus* | RASA8 | - cps |
|  | CO1122 | + cps |

### Cell wall polysaccharide competition whole cell ELISA

Cell wall polysaccharide (CWPS) competition ELISAs were carried out with Omnisera (1:1000) diluted in ELISA dilution buffer and incubated at 37°C for 30 minutes with 100 μg/ml CWPS (Statens Serum Institut) using bacteria grown in THY to an OD580nm 0.4-6 after washing in PBS and dilution to the desired CFU / ml. Fifty μl of bacterial suspension were added per well to a flat-bottomed 96 well plate and left overnight at 4°C. Plates were washed four times with a PBS + 0.05% Tween-20 wash buffer 4 times, treated with blocking buffer (PBS + 0.05% Tween-20 + 1% BSA) at 37°C for one hour, before adding Omni serum diluted 1:1000 in dilution buffer (PBS + 0.05% Tween-20 + 1% BSA) and incubating at room temperature for two hours. Following this, plates were washed four times and secondary antibody (Goat anti Rabbit IgG HRP) added at a dilution of 1:10,000, incubated for two hours at room temperature, before again washing four times. TMB substrate (100 μl) was added per well for approximately 15 minutes in the dark before stopping the reaction with the addition of sulphuric acid. The absorbance was read at 450nm subtracting readings at 550nm.

### Neutrophil-mediated opsonophagocytosis and killing

Neutrophil phagocytosis was investigated using an established flow cytometry assay using fresh neutrophils extracted from human blood and fluorescent streptococci labelled with 6-carboxyfluorescein succinimidyl ester (FAMSE; Molecular Probes). Bacterial cultures were pre-opsonised (30 mins, 37°C) with PBS, whole human sera (NHS) or heat-inactivated human sera (30 mins, 56°C) (HI-NHS). Each reaction involved 10^5^ neutrophils and a multiplicity of infection (MOI) of 10 to 1, and a minimum of 10,000 cells were analysed by flow cytometry to identify the mean percentage of neutrophils associated with bacteria. For the neutrophil killing assays, bacterial strains previously incubated in whole or heat-inactivated human serum (30 mins, 37°C) were added to fresh human neutrophils at an MOI of 1 bacterial CFU per 200 neutrophils, incubated at 37°C for 45 minutes before plating serial dilutions onto blood agar to calculate surviving CFU by colony counts after overnight incubation at 37°C, 5% CO_2_. Neutrophil killing data were expressed as a percentage survival compared to the inoculum.

### Transmission Electron Microscopy (TEM)

Mid-log-phase *S. pneumoniae* bacteria were incubated at 37°C for 20 min in phosphate buffered saline (PBS), fixed in 1% paraformaldehyde, and prepared for electron microscopy (EM) using a ruthenium red and London resin protocol as previously described (Hyams et al., 2013, Hammerschmidt et al., 2005). Bacteria were viewed using a JEOL 1010 transmission electron microscope (100 kV), and Image J software was used to determine capsule thickness. The cross-sectional area of the whole bacterium, including and excluding the capsule, was obtained and, by assuming circularity, used to calculate the bacterial radius with or without the capsule and hence the average width of the capsule layer.

### Bacterial immobilisation onto substrates for AFM experiments

Streptococcal cells were immobilised for AFM nanomechanics utilising a previously published approach (Aguayo et al., 2015). Briefly, a 100µl droplet of poly-L-lysine (PLL) or poly-dopamine (solution of 4mg/ml dopamine hydrochloride in 10mM TRIS buffer, pH 8.0) was placed on the surface of a sterile glass slide. After 1hr incubation at room temperature, surfaces were rinsed 3 times with sterile/filtered dH_2_O, dried under N_2_ airflow, and stored at 4°C. For bacterial immobilisation, 1×10^6^ CFU/ml of bacteria were harvested by centrifugation at 13,000 rpm for 10min, washed and re-suspended into 1ml of PBS to remove growth media components. Bacteria were then incubated on the coated glass slide for 10 mins, washed with PBS to remove unattached cells, and re-suspended into fresh PBS for experiments.

### Atomic Force Microscopy (AFM) experiments

All AFM nanoindentation experiments were performed on a JPK Nanowizard mounted on an Olympus IX71 inverted optical microscope (Olympus, Tokyo, Japan). MSNL-10 cantilevers (tip E, Bruker, USA) with a calculated spring constant (k) of ∼0.1 N/m, were employed throughout the study. Force mapping mode was used to obtain 16×16 pixel force maps (256 force curves per field) with a surface delay of 0 seconds and a peak force set-point of 3nN, with a constant speed rate of 2.0μm/s.

### AFM data analysis

All obtained force-curves were analysed using the JPK Data Processing Software v.5.1.8. Elasticity values (MPa) were determined by calculating the Young’s modulus according to the Derjaguin, Muller and Toporov (DMT) model. Values for maximum adhesion force (nN) and overall adhesion work (fJ) were obtained from resulting force-curves. Statistical significance was determined with the Mann-Whitney test with multiple group comparisons (p<0.05).

### Statistics

Statistical analysis was performed using GraphPad Prism 6.0. Data were presented as group means +/- the standard error of the mean (SEM). Results expressed as means were compared using either one way ANOVAs with post-hoc tests for multiple groups or students T-test when comparing the mean of two groups only. Data are representative of results obtained from at least three replicates per condition for each assay.

## Results

### Characterisation of *S. pneumoniae* TIGR4 and *S. mitis TIGR4*_*cps*_ strains using TEM

We have previously shown expression of the TIGR4 *S. pneumoniae* serotype 4 capsule fails to make the *S. mitis* SK142 strain virulent in mice (Rukke et al., 2014). To assess whether there were major morphological changes in the capsule expressed by *S. pneumoniae* TIGR4 and *S. mitis* TIGR4_*cps*_ strains, we utilised a TEM method that preserves the capsular polysaccharide to visualise and measure the depth of the capsule, whilst also observing the density and arrangement of the surface sugars. TEM demonstrated that, compared to the wild type *S. mitis* strain (Fig. 1c), which we have previously shown expresses a capsule with no known *S. pneumoniae* serotype homologue (Rukke et al., 2014), the *S. mitis TIGR4*_*cps*_ strain had a more obvious and relatively thick capsule layer (Fig. 1b) similar to the capsule layer seen with *S. pneumoniae* TIGR4 (Fig. 1a). The median thickness of the *S. mitis* TIGR4_*cps*_ capsule layer was found to be 180 nm, which is significantly lower (p<0.05) than that for *S. pneumoniae* TIGR4 (300 nm) (Fig. 1d). Quantifying capsule mass using an ELISA also demonstrated a lower quantity of serotype 4 capsule was expressed by the *S. mitis* TIGR4_*cps*_ strain compared to *S. pneumoniae* TIGR4 (Fig. 1e). Both these outcomes show that expression in *S. mitis* of the *S. pneumoniae* serotype 4 *cps* locus results in a capsule that is morphologically different to the native *S. mitis* capsule, but also in expression of a thinner capsule layer than that seen in the *S. pneumoniae* serotype 4 strain.

**Figure 1.**
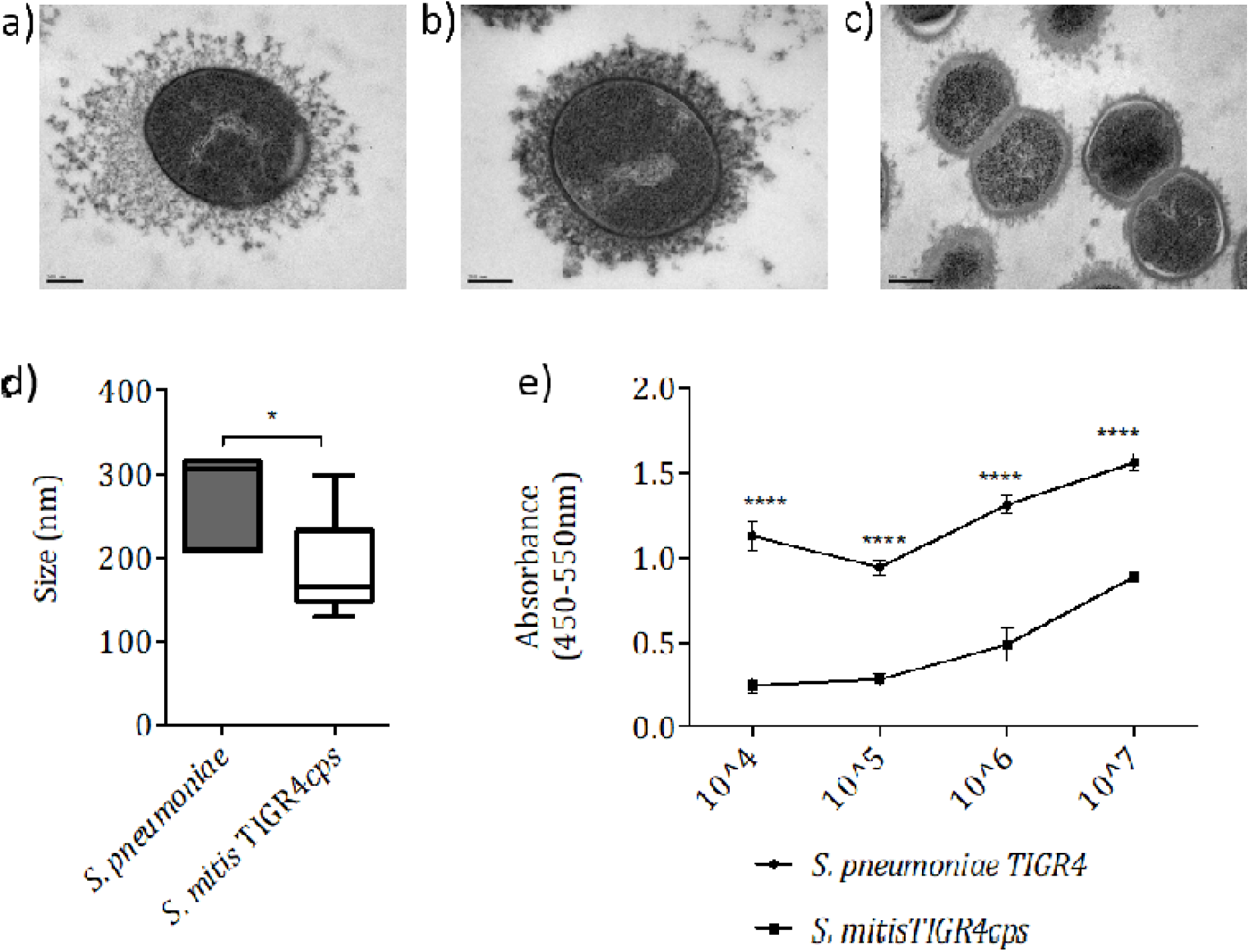
Comparing the capsule phenotype and size between wild-type *S. pneumoniae* TIGR4 and *S. mitis* expressing serotype 4. Electron microscopy images of *S. pneumoniae* TIGR4 (a) *S. mitis* TIGR4cps capsule switch strain (b) and *S. mitis* SK142. (d) Measurements from TEM images comparing capsule width of *S. pneumoniae* TIGR4 with *S. mitis* TIGR4cps mutant strain (n=5 per strain). Boxplot bars represent min/max. (e) Capsule measurement via IgG binding in Omni serum (SSI) following preincubation in 100μg cell wall polysaccharide (*** P<0.001 **** P<0.0001 two-way ANOVA with Sidak’s multiple comparisons test).

### Functional effects of expression of the *S. pneumoniae* serotype 4 capsule by *S. mitis*

Expression of the *S. pneumoniae* serotype 4 capsule by *S. mitis* reduced its sensitivity to macrophage-mediated phagocytosis (Rukke et al., 2014). To extend these data and characterise whether the serotype 4 capsule improves *S. mitis* immune evasion we used an established flow cytometry assay of neutrophil opsonophagocytosis using freshly isolated human neutrophils (Brown et al., 2002, Hyams et al., 2010b). There was a markedly higher level of neutrophil association of the *S. mitis* strain compared to the *S. mitis* TIGR4_*cps*_ and *S. pneumoniae* TIGR4 strains when opsonised in human sera (Fig. 2b). This difference was largely lost when bacteria were incubated in PBS or heat inactivated (complement inactivated) human serum (Fig, 2a, 2c). Furthermore, neutrophil killing assays showed that the high sensitivity of *S. mitis* to killing compared to *S. pneumoniae* TIGR4 was improved by expression of the serotype 4 capsule when opsonised in human serum (10% increase in survival) but not when opsonised with PBS, and with reduced differences after incubation in heat-inactivated serum (Fig. 2d). These data demonstrate that expression of the serotype 4 capsule on *S. mitis* significantly improved bacterial evasion of mainly complement-mediated phagocytosis.

**Figure 2.**
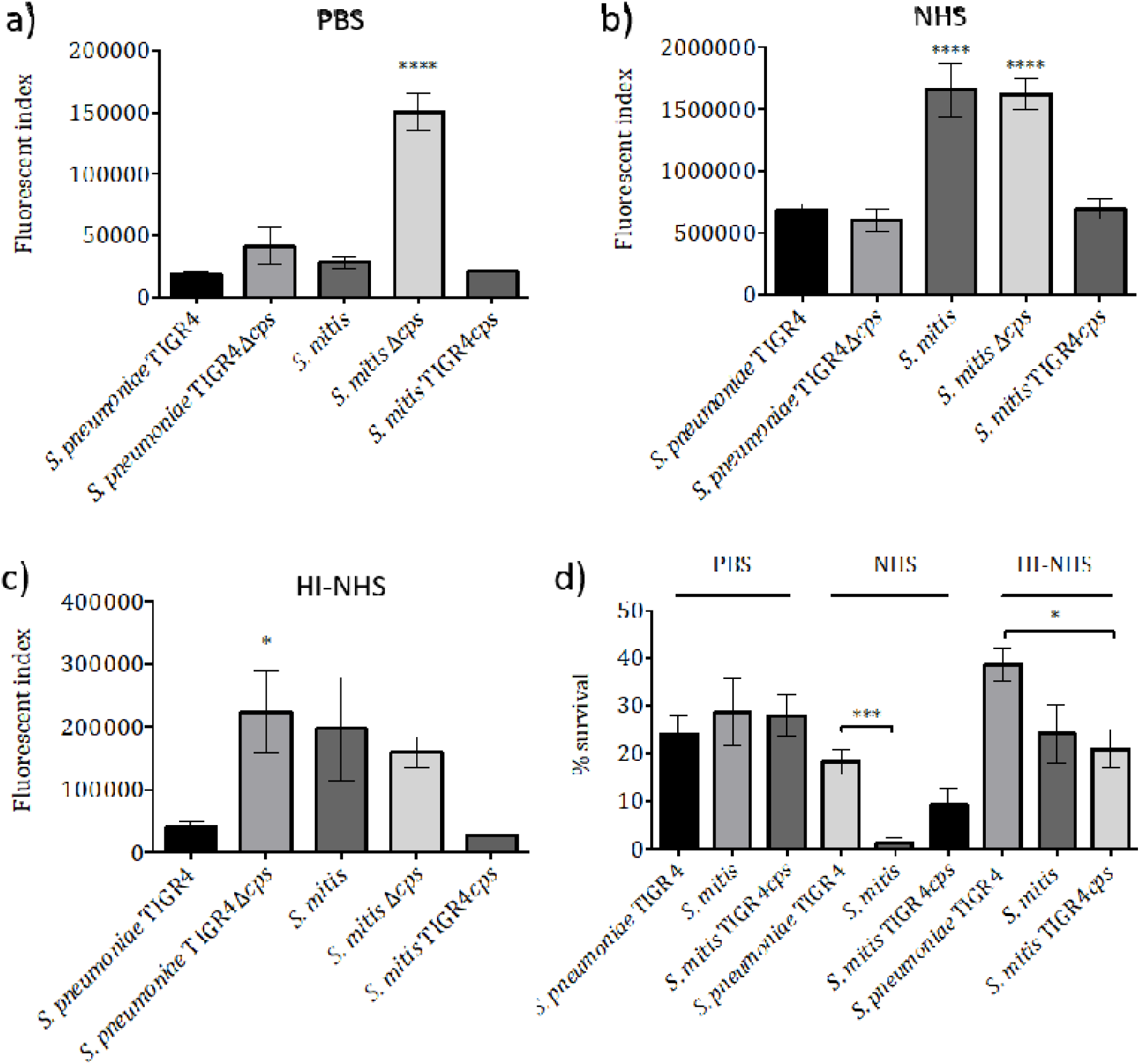
Studying the effect of the different *S. pneumoniae* and *S. mitis* capsule serotypes on neutrophil-mediated opsonophagocytosis and killing. Neutrophil uptake measured by flow cytometry following pre incubation in (A) PBS, (B) 25% whole human sera and (C) 25% heat-inactivated whole human sera (30 minutes) and incubation with human neutrophils (45 minutes). Data are presented as mean fluorescence index and error bars represent SEM. Data shown was analysed using a One-way ANOVA and a Dunnett’s multiple comparisons test comparing the mean of each column with the mean of the control column, *S. pneumoniae* TIGR4. * p<0.05 **** p<0.0001. (d) Percentage survival following pre-incubation in PBS, 25% whole human sera, (NHS) and 25% heat-inactivated whole human sera (HI-NHS) and incubation with human neutrophils at an MOI of 1:200. Error bars represent SEM. Data were analysed using an ordinary One-way ANOVA and a Dunnett’s multiple comparisons test. ** p<0.01.

### Biochemically identical capsule serotypes can differ in phenotypic appearance and size between *S. pneumoniae* and *S. mitis*

The above data demonstrate that genetic manipulation of *S. mitis* to express a *S. pneumoniae* capsular serotype results in a strain that is morphologically similar to *S. pneumoniae* with some of the protective advantages of the *S. pneumoniae* capsule compared to a native *S. mitis* capsule. Recently naturally occurring *S. mitis* strains expressing *S. pneumoniae* capsular serotypes have been identified (Skov Sørensen et al., 2016a). To assess whether the capsule expressed by these strains has a similar morphology to the same serotype expressed by *S. pneumoniae*, pairs of *S. mitis* and *S. pneumoniae* strains expressing the *S. pneumoniae* capsular serotypes 19C, 36, and 45 were investigated using TEM. Compared to *S. mitis* strains with no capsule genes contained within the site of the *cps* locus (Fig. 3g, 3h), the *S. mitis* serotype 19C, 36 and 45 strains all had a visible capsule layer (Fig. 3a-f). *S. pneumoniae* 19C and *S. mitis* SK564 *cps* locus structure are genetically identical (100% base pair identity)(Skov Sørensen et al., 2016b). TEM showed few visible differences in the capsule for both these strains - the capsule layer appeared relatively dense around the cell wall with multiple small projections giving the bacterium a slightly ‘hairy’ appearance (Fig. 3a, 3b). Using TEM to measure capsule thickness demonstrated no significant difference (Fig. 3i). Quantifying capsule mass using ELISA suggested lower levels of capsule were detected for the *S. mitis* 19C strain (Fig. 3j). The *S. mitis* SK1126 strain contains a *cps* locus with a mean nucleotide percentage identity over the 14 genes of 80-88% to the *S. pneumoniae* ST36 *cps* locus with a high sequence homology between regulatory genes, although the flippase and polymerase genes are arranged in opposite order to *S. pneumoniae* ST36 *cps* loci (Skov Sørensen et al., 2016b). Using TEM the *S. mitis* SK1126 strain capsule layer presented distinct (capsular) protrusions in the form of spikes projecting perpendicular to the cell wall (Fig. 3c, 3d). Another ST36 expressing *S. mitis* showed an almost identical phenotype profile (image not shown). Both these strains contrasted with the appearance of the *S. pneumoniae* ST36 strain capsule layer, which appeared overall denser and more homogeneous. Capsule measurements and ELISA suggest, compared to *S. mitis* SK1126, the *S. pneumoniae* strain expressing the serotype 36 capsule had a significantly thicker capsule layer (70 and 90nm respectively) and higher capsule mass (Fig. 3k, 3l). The *cps* loci of the *S. pneumoniae* ST45 and *S. mitis* SK575 strains are identical except for a short fragment of a putative acetyltransferase gene and a putative IS1381 transposase and the presence of a putative UDP-galactopyranose mutase gene just upstream of the *aliA* gene in the *S. mitis* SK575 strain (Skov Sørensen et al., 2016b). Despite this genetic similarity, *S. pneumoniae* ST45 and *S. mitis* SK575 show the clearest differences in capsule morphology by TEM. The *S. pneumoniae* ST45 capsule is dense and covers completely the cell surface, whereas the *S. mitis SK575* capsule is more sparsely distributed across the cell surface, with multiple gaps between what appear to be regularly spaced lines perpendicular to the cell wall. These lines could possibly represent capsule strands (Fig. 3e, 3f). Capsule thickness measurements and mass-quantification through ELISA suggest the *S. pneumoniae* ST45 strain capsule was significantly thicker and present a higher capsule mass when compared to the capsule of *S. mitis* SK575 (Fig. 3m, 3n).

**Figure 3.**
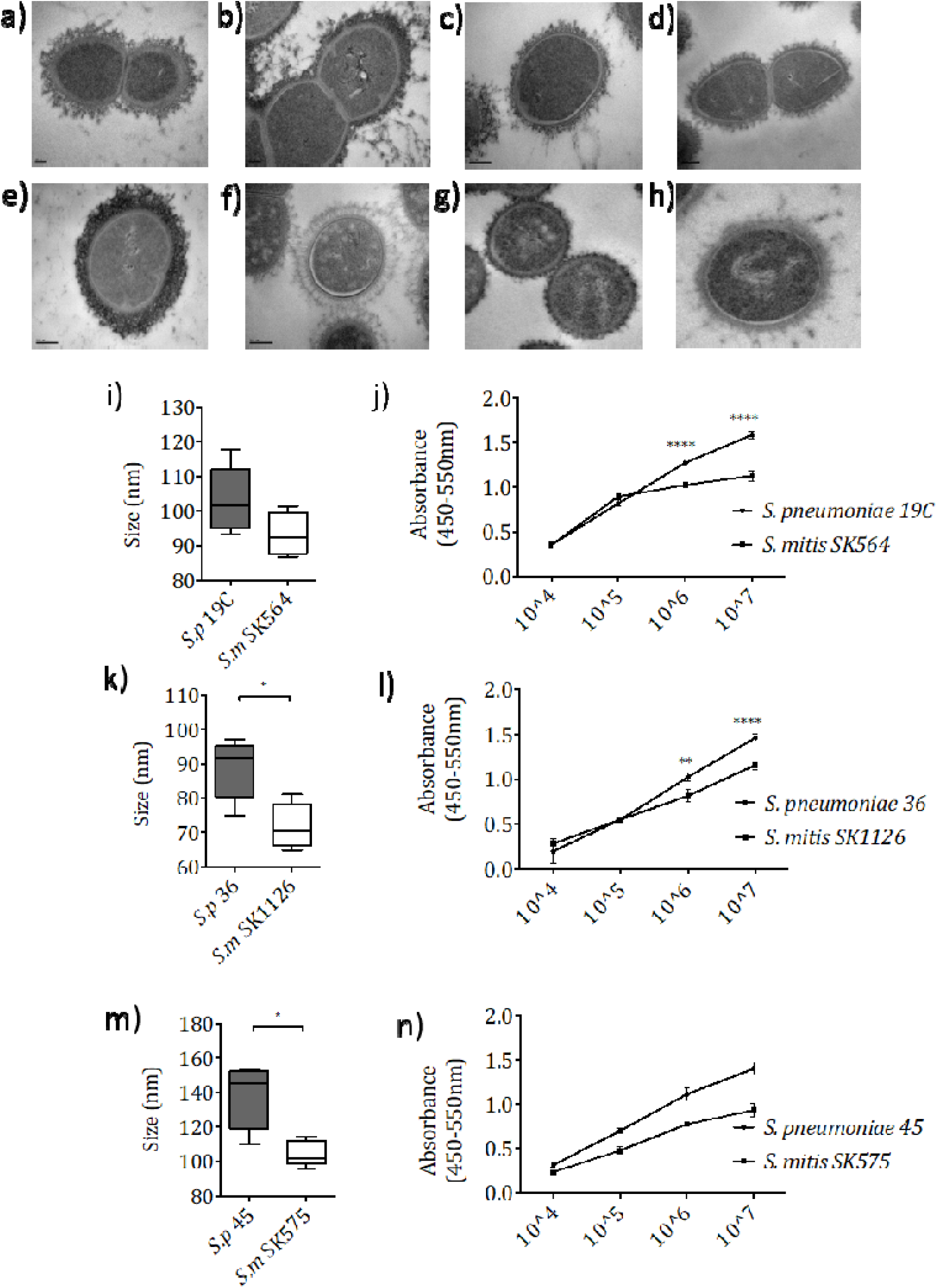
Comparing the capsule phenotype and size between strains of *S. pneumoniae* and *S. mitis* expressing matching serotypes. Electron microscopy images of (a) *S. pneumoniae* ST19C (b) *S. mitis* SK564, (c) *S. pneumoniae* ST36, (d) *S. mitis* SK1126, (e) *S. pneumoniae* ST45 (f) *S. mitis* SK575, (g) *S. mitis* SK113 (unencapsulated) and (h) *S. mitis* SK1080 (unencapsulated) when prepared using the method of LRR fixation. Scale bar shown represents 200nm in size. (i, k, m) Measurements from TEM images comparing capsule width between *S. pneumoniae* and *S. mitis* strains (n=5) Boxplots bars represent min/max. (** P<0.01 **** P<0.0001 Unpaired t-test). (j, l, n) Capsule measurement via IgG binding in Omni serum (SSI) following preincubation in 100μg cell wall polysaccharide (*** P<0.001 **** P<0.0001 two-way ANOVA with Sidak’s multiple comparisons test).

### Comparative sensitivity to neutrophil-mediated killing of *S. mitis* and *S. pneumoniae* strains naturally expressing same capsular serotype

The susceptibility to neutrophil phagocytosis of the *S. mitis* and *S. pneumoniae* strains expressing the *S. pneumoniae* capsular serotypes 19C, 36, and 45 was assessed using a neutrophil killing assay after opsonisation in PBS or 25% baby rabbit complement (Hyams et al., 2010a). When pre-opsonised with PBS, all *S. pneumoniae* strains tested displayed a 2- to 5-fold increase in CFU over time, representing bacterial replication and significant resistance to neutrophil killing (Fig. 4a). The results for the *S. mitis* strains were not as well defined as *S. pneumoniae, S. mitis* SK1126 showed a similar increase in CFU as the corresponding capsular serotype 36 *S. pneumoniae* strain, whereas the *S. mitis* ST19C and ST45 strains both had very poor survival even when opsonised with PBS. When opsonised with complement (25% BRC) there was a marked reduction in CFU recovered for the *S. mitis* SK1126 strain whereas the capsular serotype 36 *S. pneumoniae* strain was still resistant to neutrophil killing. Complement improved neutrophil mediated killing of the *S. pneumoniae* ST19C and ST45 strains, and reduced the differences to the *S. mitis* ST19C and ST45 strains that were seen when the bacteria were opsonised with just PBS (Fig. 4b). Overall, the *S mitis* strains showed greater sensitivity to neutrophil killing than the *S. pneumoniae* strains expressing the same capsule, although for the *S. mitis* SK1126 strain (but not the *S. mitis* ST19C and ST45 strains) this required complement to be detectable.

**Figure 4.**
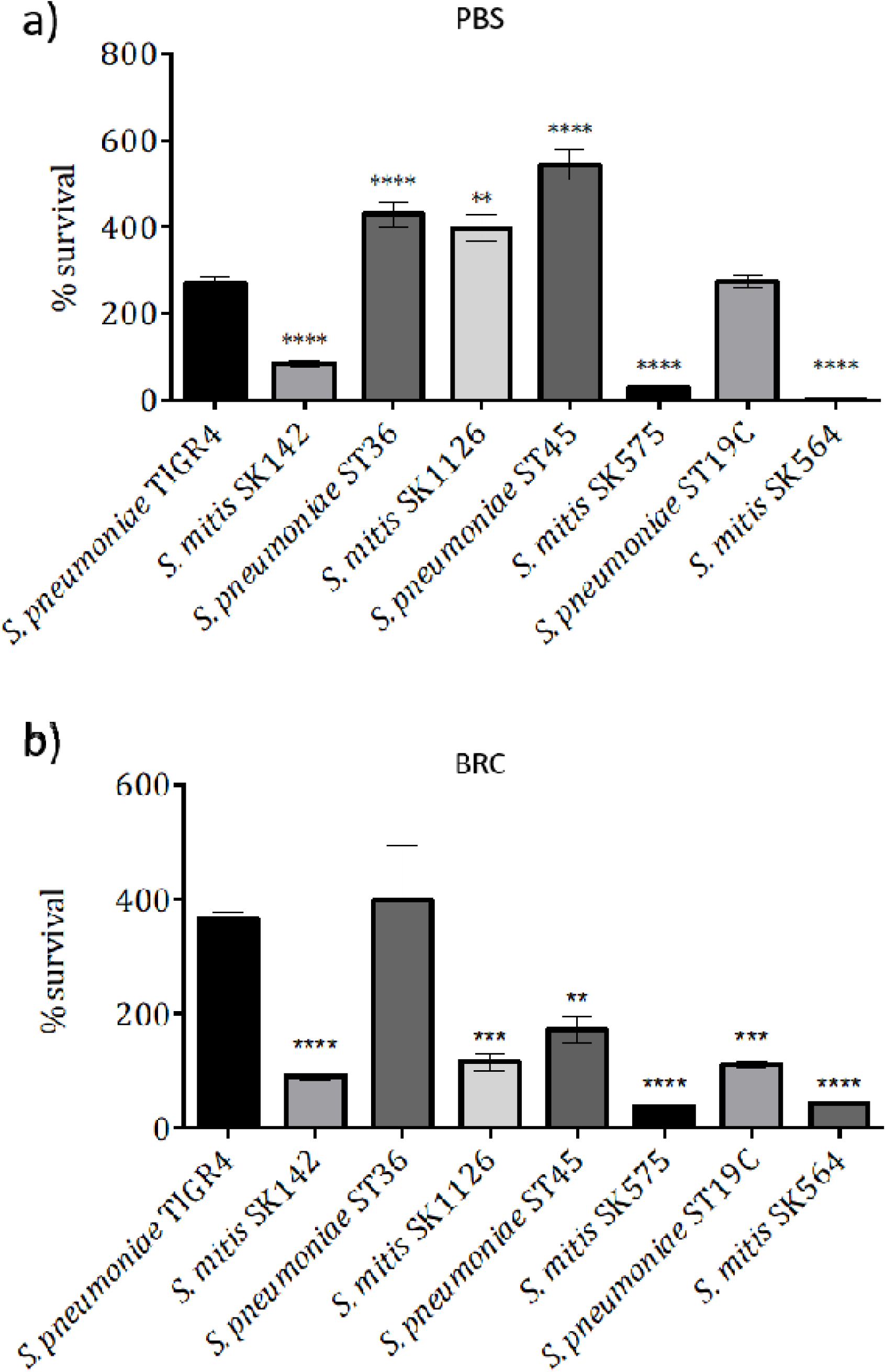
Investigating the effect of different naturally occurring *S. mitis* capsule serotypes on neutrophil-mediated killing. Percentage survival following pre-opsonisation in (A) PBS, (B) 25% baby rabbit complement before incubation with human neutrophils at an MOI of 1:200 for 45 minutes. Error bars shown represent SEM. Survival of less than 100 represents bacterial killing, whereas % survival greater than 100 represents bacterial growth. Data were analysed using a One-way ANOVA and a Dunn’s multiple comparisons test comparing the mean of each column with the mean of the control column. Control columns were *S. pneumoniae* TIGR4 and *S. mitis* SK142. * p<0.05 ** p<0.01 *** p<0.001 **** p<0.0001.

### Mechanical stiffness of streptococcal capsules

To investigate the mechanical properties of streptococcal capsules, we initially used AFM-based indentation to map the elasticity of the *S. pneumoniae* TIGR4, *S. mitis* TIGR4_*cps*_, *S. mitis* SK142 strains and their unencapsulated derivatives. To evaluate the properties of the capsules specifically, measurements were acquired both in the centre of the cell (body) and as close to the edge of the cell (edge) as possible. By performing these differential measurements, it became possible to decouple the mechanical response of the complex cell-capsule at the centre (body) and capsule only (edge). Indentation measurements performed directly over the cell body of *S. pneumoniae* TIGR4, *S. mitis* TIGR4_*cps*_, and *S. mitis* SK142 strains and their unencapsulated derivatives yielded little to no differences between the approach or retraction curves suggesting a very good mechanical compliance of the bacterial cell, acting as an almost perfect elastic material (Table 2). Additionally, little to no adhesion could be recorded over the cell body regardless of the presence or absence of capsules. It is worth noting that if both the indentation load and the cantilever’s spring constant were lower (load <0.5nN and k <0.05N/m) a much stronger adhesion response could be recorded (data not shown). Overall, the body of these bacterial strains’ cells presented a very elastic and compliant behaviour with values ranging from 0.87 MPa for *S. mitis* TIGR4_*cps*_ to 9.14 MPa for *S. mitis* Δ*cps*.

**Table 2:** Elasticity, adhesion forces and energy for *Streptococcus pneumoniae* and *Streptococcus mitis* strains. Data is presented as medians (IQR).

| | *S. pneumoniae*<br>TIGR4 | *S. pneumoniae*<br>TIGR4 $\Delta cps$ | *S. mitis*<br>SK142 | *S. mitis*<br>SK142 $\Delta cps$ | *S. mitis*<br>SK142<br>TIGR4*cps* |
| --- | --- | --- | --- | --- | --- |
| **Elasticity body**<br>**(MPa)** | 1.79<br>(0.38-2.51) | 1.95<br>(1.17-3.13) | 6.09<br>(2.94-9.72) | 9.14<br>(0.30-33.28) | 0.87<br>(0.14-3.21) |
| **Elasticity edge**<br>**(MPa)** | 0.32<br>(0.17-0.55) | --- | 2.63<br>(0.89-4.17) | --- | 0.03<br>(0-0.66) |
| **Adhesion**<br>**(nN)** | 0.17<br>(0.09-0.29) | 0.11<br>(0.07-0.18) | 0.40<br>(0.27-0.54) | 0.16<br>(0.06-0.27) | 0.06<br>(0.03-0.10) |
| **Energy**<br>**(fJ)** | 0.01<br>(0-0.08) | 0.01<br>(0-0.01) | 0.05<br>(0.01-0.13) | 0.02<br>(0-0.11) | 0<br>(0-0.03) |

Performing measurements on the cell edge for the encapsulated strains (*S. pneumoniae* TIGR4, *S. mitis* TIGR4_*cps*_, *S. mitis* SK142 strains) presented a different indentation pattern. A hysteresis between the approach and retraction curves together with non-specific adhesion (integrated area under the retraction curve) could be recorded in the presence of the capsule at the edge of the bacteria, where we are only indenting the capsule (Fig. 5a, 5c). The hysteresis between the approach and retraction curve implied a significant change in the mechanical properties of the sample between the indentation and the relaxation of the material i.e. this region of the sample displays a visco-elastic behaviour (Efremov et al., 2017), due for example to a reorganisation of the capsule following the initial indentation. Viscoelasticity is revealed in a clear hysteresis between the approach and retraction parts of curves (Rebelo et al., 2013). In terms of elastic modulus, the edge of *S. pneumoniae* TIGR4 (E= 0.32 MPa) showed a significant decrease (∼1.5 MPa) compared to the body (E= 1.79 MPa). In the case of the isogenic capsule deletion mutant (Fig. 5b), measurements taken for the cell body showed a minor increase in elastic modulus (E=1.95 MPa). Overall, these findings indicate that the cell body of *S. pneumoniae* TIGR4 and *S. pneumoniae* TIGR4 Δ*cps* is significantly stiffer than the cell edge. Furthermore, a clear change in the biomechanical properties of the bacterial cell can be expected when the surface capsule is lost.

**Figure 5.**
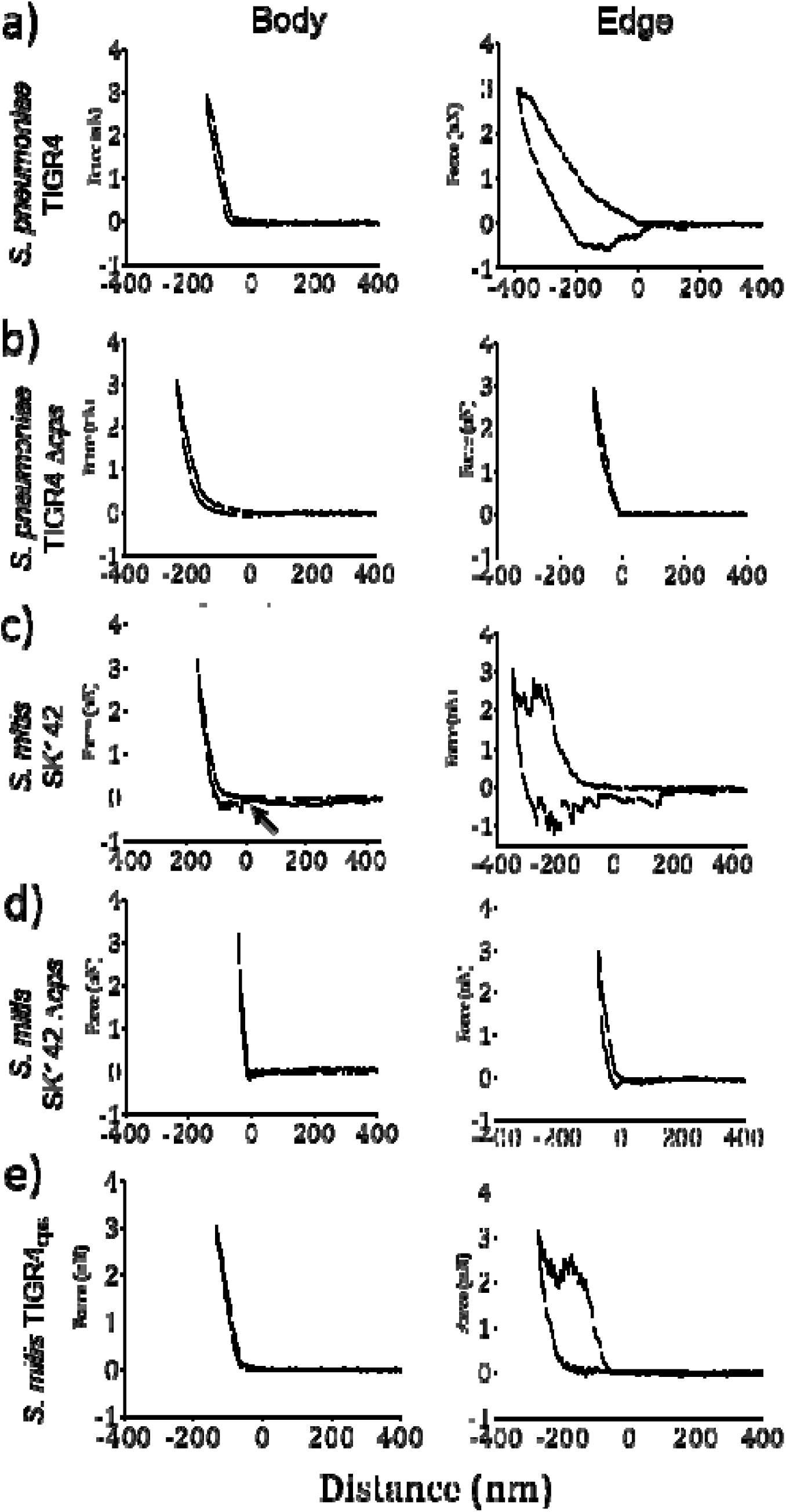
Studying the nanoscale elastic properties of Streptococcal capsular material. AFM was used to obtain force-curves on both the cell centre and edge, in order to obtain information on the elastic and adhesive properties of each bacterium. Representative force curves are shown for each area. (c) Histograms corresponding to the elasticity observed on the central area of the cell body of each strain are shown. Non-linear gaussian fits are shown. Inset of *S. pneumoniae* TIGR4 represents the histogram for the left peak (lower values).

Similar to *S. pneumoniae*, the mechanical response of *S. mitis* (SK142) presented some hysteresis between the approach and retraction curves on the cell edge together with a significant amount of adhesion (Fig. 5c). The elastic modulus of cell body median of *S. mitis* (E= 6.09 MPa) was over 3 MPa stiffer than the cell edge (E= 2.63 MPa). In the indentation measurement performed on the *S. mitis* wild type (SK142) cell body, the retraction curve also presented specific unbinding events. Of all the strains examined in this study, these events were seen only with wild-type *S. mitis* strains. In comparison to *S. pneumoniae* TIGR4, the cell body modulus of elasticity of *S. mitis* was over 4 MPa higher (E= 1.79 and 6.09 MPa respectively), though the differences in moduli of elasticity for *S. pneumoniae* TIGR4 and *S. mitis* SK142 edges were less distinct than that of the cell body (E= 0.32 and 2.63 MPa respectively). The *S. mitis* Δ*cps* strain demonstrated similar biomechanical characteristics to the *S. pneumoniae* TIGR4 Δ*cps* strain with similar approach and retraction curves (Fig. 5d), and a sharp gradient from point of contact to set loading force for both the bacterial cell body and the bacterial edge. The unbinding events seen for *S. mitis* wild type were not observed in the case of *S. mitis* Δ*cps*. Finally, the median elasticity values for the cell body of *S. mitis* Δ*cps* demonstrated an increased spread of values of elastic modulus, implying a much greater mechanical heterogeneity of this cell when compared to any of the other strains investigated in this study.

To determine whether the capsule serotypes considered identical by biochemistry would also possess the same biomechanical properties when expressed in *S. mitis*, indentation measurements were also carried out on the *S. mitis* TIGR4_*cps*_ strain. The indentation curves acquired directly on the cell body again of the *S. mitis* TIGR4_*cps*_ strain showed a similar approach and retraction curves to *S. mitis* (Fig. 5e). However, the median elastic modulus for the cell body of *S. mitis* TIGR4_*cps*_ was significantly lower than that of the *S. mitis* wild-type strain (E= 0.87 and 6.09 MPa respectively), bringing it much closer to that measured for the *S. pneumoniae* TIGR4 cell body (E= 1.79 MPa). As shown for the encapsulated TIGR4 and *S .mitis* SK142 wild type strains, the curves obtained from the cell edge also presented the previously observed hysteresis pattern between the approach and the retraction curve (Fig. 5e). Finally, in contrast to *S. pneumoniae* TIGR4, the median elasticity value for the cell edge of *S. mitis* TIGR4_*cps*_ was extremely small (E= 0.03 MPa) suggesting it was much softer when compared to the other strains.

### *S. pneumoniae* vs. *S. mitis*: Contribution of the surface capsule to adhesion

Adhesion force (between the AFM probe and individual bacterial cell) was calculated for each strain from the retraction curve of the force maps. Interestingly, both strains of bacteria without any capsule (Fig. 6 b, d) presented the same distribution of adhesion with very low median values suggesting that the bacteria were not adhering very well to the cantilever. When both these strains were functionalised with the TIGR4 capsule, they presented again a similar pattern distribution of adhesion to one another (albeit with higher values that in the no-capsules case) as presented in Fig 6 a & e. It appears that the presence of the TIGR4 capsule led to an increase median of adhesion for both these strains to 0.18 nN. Finally, the strongest adhesion behaviour was observed when *S. mitis* was functionalised with SK142 capsule (Fig. 6c). In this case, the adhesion increases significantly to reach a median of 0.40 nN when compared to the capsule deletion or TIGR4_*cps*_. This approach strongly suggests that it capsular serotype can play a direct part in the strength of bacterial adhesion to external surfaces.

**Figure 6.**
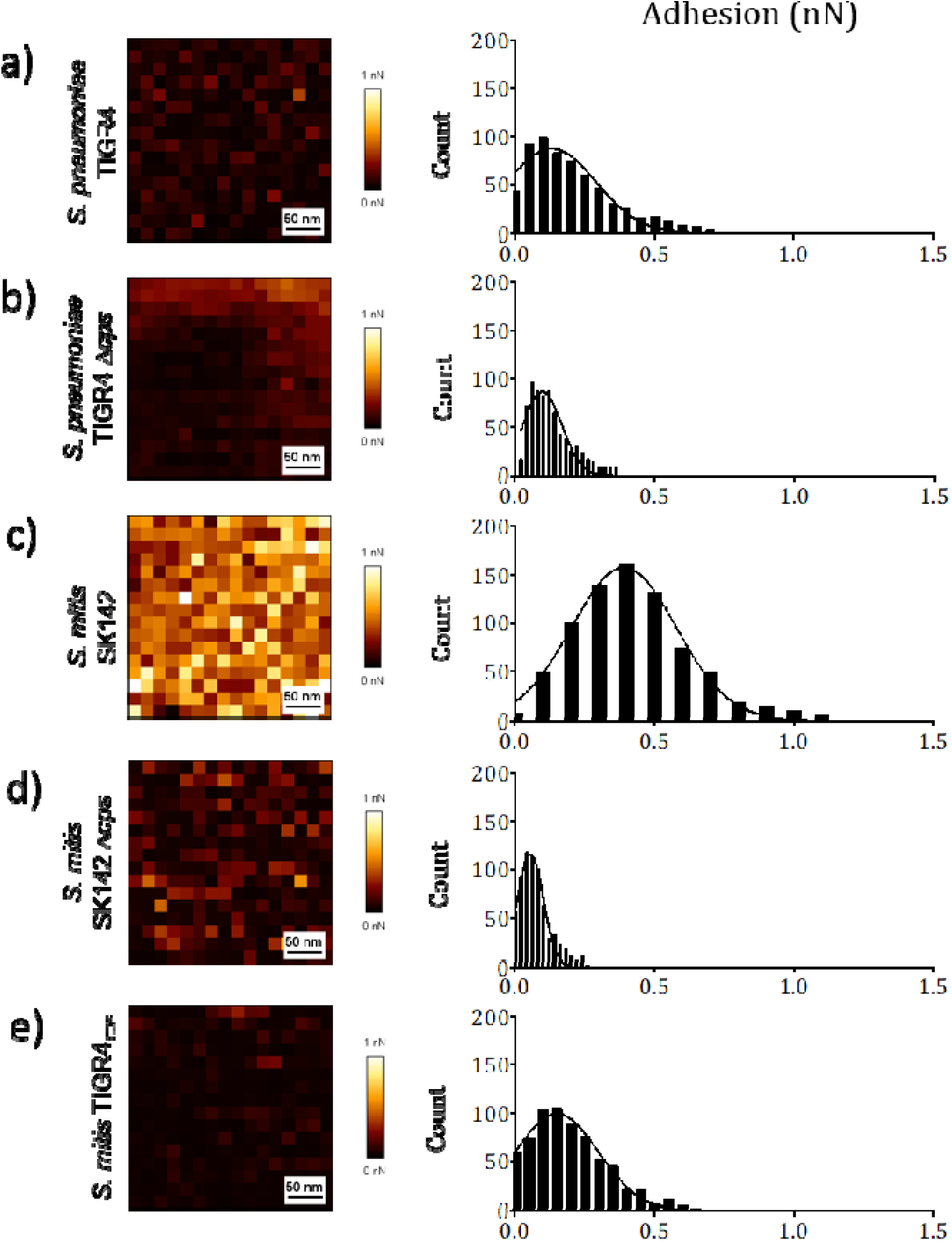
Capsule of *S. pneumoniae* and *S. mitis* is important for strain adhesive properties. FVI maps representing 300×300nm areas on the centre of the cell body and histograms for adhesive forces observed for (a) *S. pneumoniae* TIGR4, (b) *S. pneumoniae* TIGR4 Δ*cps*, (c) *S. mitis* SK142, (d) *S. mitis* Δ*cps* and (e) *S. mitis* TIGR4_cps_.

### Mechanical properties of bacterial edges for encapsulated and unencapsulated *S. aureus* strains confirm the AFM data reflect capsule effects

To further characterise the effect of capsule loss on the nanomechanical properties of bacteria, *S. aureus* RASA8 and its corresponding unencapsulated strain CO1122 (Spoor et al., 2015) were explored structurally by both TEM and mechanically by AFM indentation. TEM imaging was carried out to confirm the presence and absence of capsule in both strains, respectively (Fig. 7a, 7c). Similar to the observations with streptococci, force-curves observed with the encapsulated strain demonstrated an increased hysteresis in the force-distance curve as well as an increased adhesion work in comparison to the unencapsulated strain (Fig. 7b, 7d), characteristic of increased adhesion between the microbial cell and surface, and indeed the adhesion force measured for *S. aureus* RASA8 was higher than that for *S. aureus* CO1122 (0.42 and 0.15 nN respectively) (Fig. 7f, 7g, 7h). Similar to the pattern of elastic modulus observed for *S. mitis* SK142 and its corresponding unencapsulated strain, *S. aureus* CO1122 demonstrated a significantly increased cell body elastic modulus to that of *S. aureus* RASA8 (E= 13.36 and 4.18 MPa respectively) (Fig 7e). RASA8 also demonstrated different elastic moduli values for both the cell centre and cell edge, consistent with the presence of capsular material surrounding the bacterium. The same behaviour was not observed for the unencapsulated CO1122 strain, suggesting it directly reflects the mechanical properties of the polysaccharide capsule. Overall, loss of capsule was found to display a similar AFM pattern for both streptococci and staphylococci strains utilised in this study, confirming the data obtained is related to capsule effects on the physical properties of the bacterial surface.

**Figure 7.**
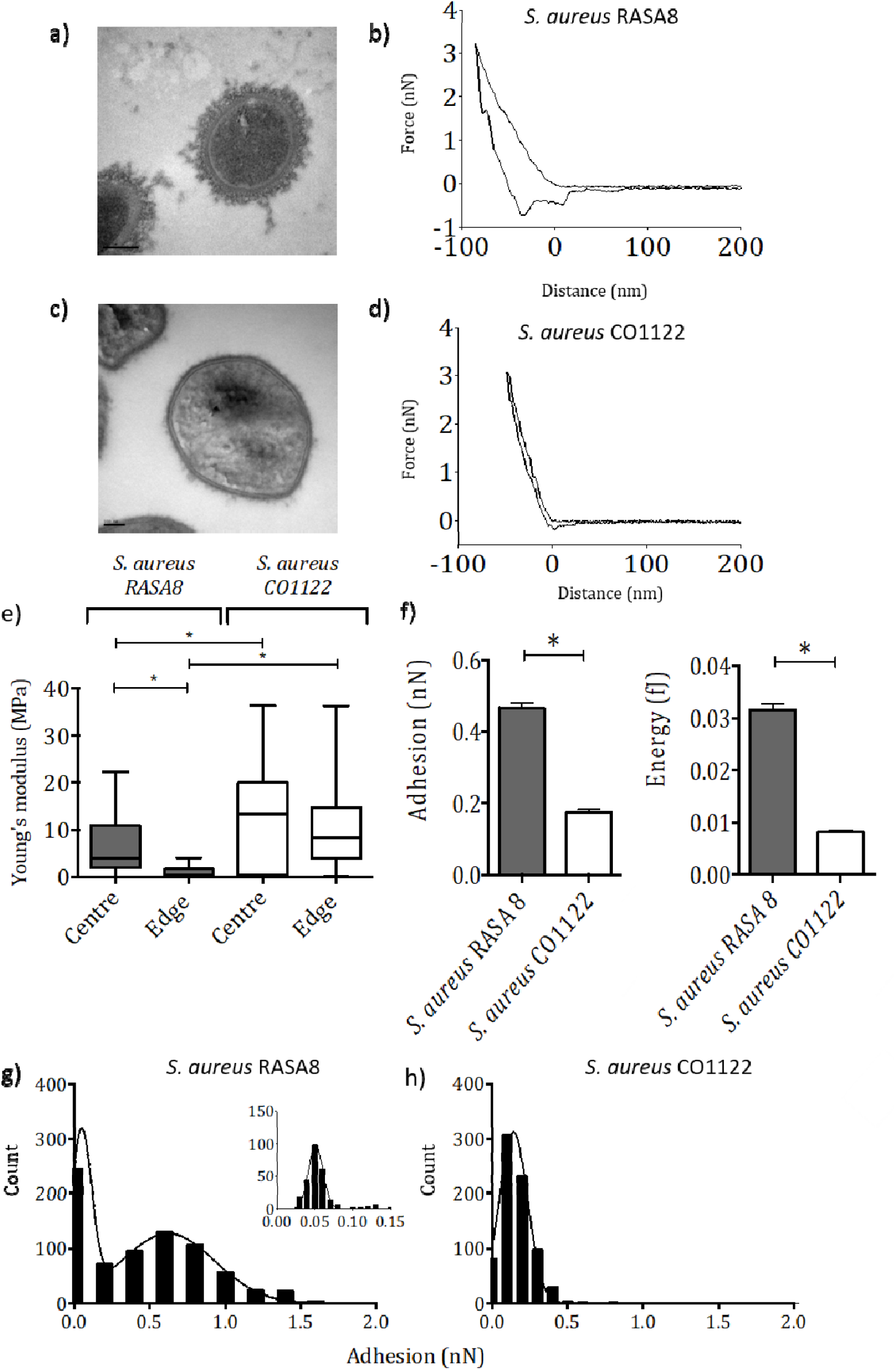
Validating the described protocol on *S. aureus* strains. Electron microscopy and force curves obtained from FVI maps on the surface of (a) and (b) *S. aureus* RASA8 (encapsulated) and (c) and (d) *S. aureus* CO1122 (unencapsulated). (e) Box plots of elasticity on cell centre and edge for both strains. Boxplot bars represent median and IQR. (d) and (e) represent data on adhesion recorded at the centre of each bacterium surface. Data were analysed using a Kruskal-Wallis One-way ANOVA. * p<0.05

### Comparison of the biophysical properties of the ST36 *S. pneumoniae* and *S. mitis* capsules

Both *S. pneumoniae* ST36 and *S. mitis* SK1126 naturally express a biochemically identical serotype 36 capsule polysaccharide, but EM and ELISA data demonstrated differences in the physical structure and mass of the capsule (Fig. 3c, 3d, 3k, 3l) with the *S. pneumoniae* strain showing markedly more resistance to complement mediated neutrophil killing (Fig. 4b). AFM imaging (Fig. 8e, 8k) confirmed that these encapsulated strains consisted of an oval body surrounded by an edge of material that likely represents the capsule, and thereby supports the concept that the force-distance data from the bacterial edge were due to the capsule alone. Force mapping was carried out on these strains to determine if the species background affected the biophysical characteristics of the capsule serotype produced. Force-distance curves obtained from the *S. pneumoniae* ST36 cell body exhibited a steep gradient and no separation of the approach and retraction curve (Fig. 8a). Force-distance curves generated from the cell edge showed a distinct bimodal pattern of a steep incline to between approximately 2–2.5nN force, after which the force dropped down to almost 1nN before being reapplied by the AFM and 3nN reached (Fig. 8b). This relaxation in the approach curve of the force-distance cycle suggests that the material being indented (in this case the capsule) is being perforated by the AFM probe. As the maximum indentation load is not reached, the force-distance cycle continues. The steep second part of the approach would correspond to the indentation of the body of the cell. The overall elasticity of the *S. pneumoniae* ST36 cell body was approximately 2 MPa higher than the cell edge; both cell body and edge demonstrated a skewed frequency distribution towards the lower MPa scale (Fig. 8c, 8d). A similar bimodal force curve profile was exhibited by *S. mitis* SK1126, showing a steep curve with no hysteresis for the cell body (Fig. 8g), whereas the frequency distribution of the cell edge was shifted to the left (Fig. 8i). The *S. mitis* SK1126 overall body elasticity was approximately 3 MPa greater than the cell edge (Fig. 8h, 8j). Distribution of adhesion values for both serotype 36 expressing capsule strains demonstrated almost identical distribution from 0.08 – 0.15nN. A small increase in the median adhesion force of *S. mitis* SK1126 (0.12 nN) was seen in comparison to *S. pneumoniae* ST36 (0.1 nN), but there was no difference in energy (adhesion work) between the two strains (Fig. 8f, 8l). These data along with calculated elasticity demonstrate that compatible with their identical biochemical structure, the serotype 36 capsule layers of *S. pneumoniae* ST36 and *S. mitis* SK1126 have similar biomechanical characteristics which have a distinct pattern when compared to those for the serotype 4 capsule.

**Figure 8.**
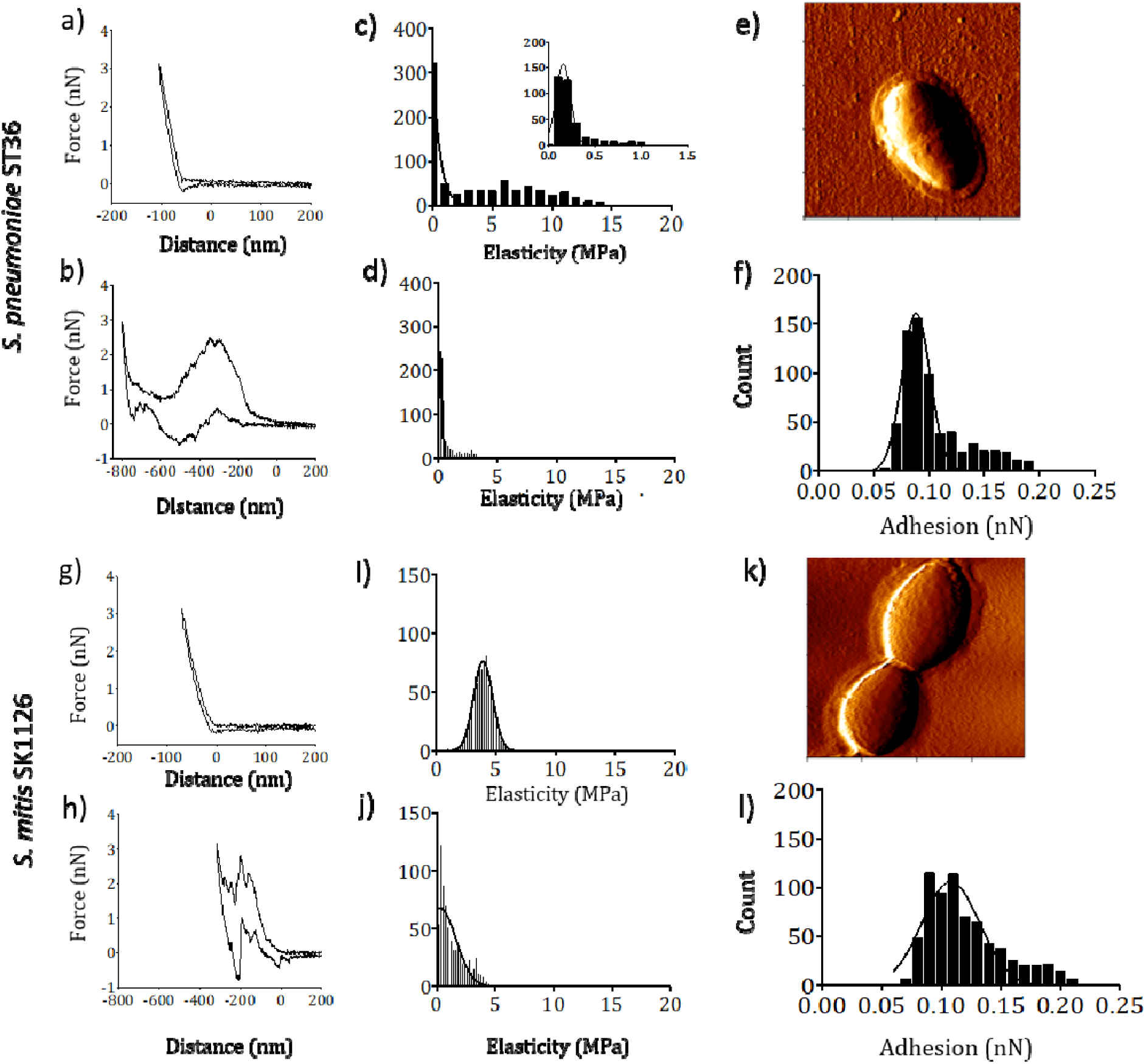
Comparison between the mechanical properties of different naturally occurring serotype 36 strains. (a) and (c) correspond to force curves obtained at the centre and edge of *S. mitis* SK1126, respectively. (b) and (d) Elasticity histograms for both centre and edge of *S. mitis* SK1126. (e) AFM imaging of *S. mitis* SK1126. (f) Histogram of adhesion forces recorded for the centre of *S. mitis* SK1126 cells. (g) and (i) correspond to force curves obtained at the centre and edge of *S. pneumoniae* ST36, respectively. (h) and (j) Elasticity histograms for both centre and edge of *S. pneumoniae* ST36 (k) AFM imaging of *S. pneumoniae* ST36. (l) Histogram of adhesion forces recorded for the centre of *S. pneumoniae* ST36 cells.

## Discussion

The bacterial capsule is a well-recognised major virulence determinant for *S. pneumoniae.* However, *S. mitis*, its low pathogenic close genetic relative often also expresses a surface capsule as further confirmed by our TEM and AFM images (Fig. 1, 3, 8). However, differences were observed between the morphological and biomechanical properties of the *S. pneumoniae* capsular serotypes expressed in *S. mitis* strains compared to expression in *S. pneumoniae*. For example, capsule width was significantly diminished in *S. mitis* compared to *S. pneumoniae* (Fig. 3i, 3k, 3m). Furthermore, capsules of *S. pneumoniae* ST36 and *S. mitis* SK1126, which are biochemically identical, were found to have different capsule mass and morphological structure (Fig. 3c, 3d, 3k, 3l). These data suggest that although the influence of genes on the chemical makeup of *S. pneumoniae* capsule serotype has been partially determined (Bentley et al., 2006), other unknown factors also influence the potential capsule morphology and biophysical properties. Potential possibilities for these factors include differences in the supply of capsular monosaccharide units, capsule assembly or regulators of capsule size. For example, allelic variation of the Spn556II type-I restriction modification system is associated with phase variation of *S. pneumoniae* between transparent (thin capsule) and opaque (thick capsule) phases (Li et al., 2016, Manso et al., 2014), but this system is not present in *S. mitis* (Skov Sørensen et al., 2016a). Our results demonstrate that *S. mitis* strains show an increased susceptibility to neutrophil killing *in vitro*, when compared to *S. pneumoniae* strains naturally expressing the same capsule serotype. Thus, despite expression of identical serotypes, the biological capabilities conferred to *S. mitis* and *S. pneumoniae* are different, which may play a role on their virulence *in vivo*. This may be explained by the reduced capsule width observed for the *S. mitis* strains compared to *S. pneumoniae*, confounded with other genetic changes. Further investigation of a wider range of strains and serotypes could shed further light on this association.

Previous reports in literature have demonstrated the use of AFM to image and characterise the elastic and adhesive properties of bacterial cells (Dufrêne, 2014, Su et al., 2012, Wang et al., 2013, Wang et al., 2015). In this work, we have utilised a non-invasive bacteria immobilisation approach that allowed the imaging and probing of *S. mitis* and *S. pneumoniae* attached to biopolymer-coated glass substrates, in PBS. We observed distinct force-curve profiles and elastic modulus data for the centre and edge areas of each bacterium for *S. pneumoniae* TIGR4, *S. mitis* and *S. mitis* TIGR4_*cps*_. All encapsulated strains were found to possess two distinct areas of elastic properties, namely, a “stiffer” centre surrounded by a “softer” edge (Table 2). This “edge effect” was not observed in the non-encapsulated strains, and therefore we believe it is a direct representation of the presence of bacterial capsule. This model was further confirmed by utilising *S. aureus* RASA8 and its corresponding unencapsulated strain CO1122, which also demonstrated elastic and adhesive differences associated with the presence of capsule (Fig. 7).

The unbinding profiles observed for the capsule of *S. pneumoniae* TIGR4, *S. mitis* and *S. mitis* TIGR4_*cps*_ demonstrated the characteristic saw-tooth like patterns of single-molecule unbinding (Strunz et al., 1999), and thus suggest the interaction of single adhesive units between the capsule and AFM probe. Utilising this approach, we were also able to demonstrate similar nanomechanical properties and unbinding patterns for two serotype 36 capsule strains, *S. pneumoniae* ST36 and *S. mitis* SK1126. Furthermore, the force-curve profiles for capsular ST36 strains were markedly different from the serotype 4 capsule (Fig. 5a, 5e and 8b, 8h). Our AFM results also show increased adhesion forces for *S. mitis* compared to *S. pneumoniae* TIGR4, *S. mitis* TIGR4_*cps*_, and their unencapsulated derivatives. Unencapsulated strains demonstrated the lowest adhesion values, which supports data showing the capsule impairs the adhesion of streptococci to cell surfaces (Hammerschmidt et al., 2005, Magee and Yother, 2001, Nelson et al., 2007). Differences amongst the biophysical properties of *S. pneumoniae* TIGR4 and *S. mitis TIGR4*_*cps*_ were observed, compatible with the data showing morphological disparities observed with TEM between these strains (Fig. 1a, 1b). Significant differences were also found between encapsulated strains, with different capsule serotypes having distinct adhesion profiles. The data on neutrophil sensitivity, EM findings, and biophysical characteristics measured using AFM for strains expressing difference capsular serotypes are summarised in Table 3. Overall, our results demonstrate that the AFM analysis of streptococcal capsular material can characterise differences at the nanoscale in the physical properties of the capsule between bacterial species and strains.

**Table 3:** Summary table of the physical characteristics of the capsule and neutrophil killing sensitivity for streptococcal strains investigated using AFM and EM.

| Strain | ST* | Mean capsule width (nm) | Neutrophil killing sensitivity | Body elasticity (MPa) | Capsule elasticity (MPa) | Adhesion force (nN) |
| --- | --- | --- | --- | --- | --- | --- |
| *S. pneumoniae*TIGR4 | 4 | 270 | + | 1.79 | 0.32 | 0.17 |
| *S. mitis* SK142 TIGR4cps | 4 | 185 | ++ | 0.87 | 0.03 | 0.06 |
| *S. mitis* SK142 wild type | novel | 45 | ++++ | 6.09 | 2.63 | 0.4 |
| *S. pneumoniae* ST36 | 36 | 88 | + | 3.713 | 0.84 | 0.1 |
| *S. mitis* SK1126 | 36 | 72 | ++++ | 3.224 | 1.1 | 0.12 |

\*Capsular serotype

## Conclusion

Overall, although *S. mitis* was found to express the same chemical capsule serotypes as *S. pneumoniae*, there were important morphological and functional differences. *S. mitis* strains expressed a thinner capsule layer than corresponding capsular serotype *S. pneumoniae* strains, which also correlated to an increased susceptibility to neutrophil killing. Previously linking how variation in the chemical structure of the capsule influences the biophysical properties of the bacterial surface to influence bacterial immune evasion and other phenotypes was not possible. We have now shown that AFM can define the biophysical properties of the capsule of living *S. mitis* and *S. pneumoniae*, reproducibly quantifying the elastic and adhesive properties of bacterial cell surfaces. AFM was able to measure both the properties of the capsule and bacterial cell body, allowing the investigation of whether both factors combined are important for the biology of the organisms’ interactions with the host. By being able to decipher the mechanically compliant properties and adhesion of different capsules as a function of their serotypes, we have demonstrated that it is now possible to assess bacteria species as a functional biomechanical living entity. Future work will use AFM to study the biophysical phenotypes of different capsule serotypes across a larger number of strains to identify the relationship between capsule structure, its physical properties, and biological effects on virulence and surface adhesion.

## Acknowledgments

We would like to thank the London Centre for Nanotechnology (London UK) for providing access to the AFM Facility and Dr Richard Thorogate for his helpful support in terms of AFM nanometrology.

